# Potent Immunogenicity and Broad-Spectrum Protection Potential of Microneedle Array Patch-Based COVID-19 DNA Vaccine Candidates Encoding Dimeric RBD Chimera of SARS-CoV and SARS-CoV-2 Variants

**DOI:** 10.1101/2022.12.01.518127

**Authors:** Feng Fan, Xin Zhang, Zhiyu Zhang, Yuan Ding, Limei Wang, Xin Xu, Yaying Pan, Fang-Yuan Gong, Lin Jiang, Lingyu Kang, Zhihua Kou, Gan Zhao, Bin Wang, Xiao-Ming Gao

## Abstract

Breakthrough infections by SARS-CoV-2 variants pose a global challenge to pandemic control, and the development of more effective vaccines of broadspectrum protection is needed. In this study, we constructed pVAX1-based plasmids encoding heterodimeric receptor-binding domain (RBD) chimera of SARS-CoV and SARS-CoV-2 Omicron BA.1 (RBD^SARS/BA1^), SARS-CoV and SARS-CoV-2 Beta (RBD^SARS/Beta^), or Omicron BA.1 and Beta (RBD^BA1/Beta^) in secreted form. When i.m. injected in mice, RBD^SARS/BA1^ and RBD^SARS/Beta^ encoding plasmids (pAD1002 and pAD131, respectively) were by far more immunogenic than RBD^BA1/Beta^ plasmid (pAD1003). Dissolvable microneedle array patches (MAP) laden with these DNA plasmids were fabricated. All 3 resulting MAP-based vaccine candidates, namely MAP-1002, MAP1003 and MAP-131, were comparable to i.m. inoculated plasmids with electroporation assistance in eliciting strong and durable IgG responses in BALB/c and C57BL/6 mice as well as rabbits, while MAP-1002 was comparatively the most immunogenic. More importantly, MAP-1002 significantly outperformed inactivated SARS-CoV-2 virus vaccine in inducing RBD-specific IFN-γ^+^ T cells. Moreover, MAP-1002 antisera effectively neutralized pseudoviruses displaying spike proteins of SARS-CoV, prototype SARS-CoV-2 or Beta, Delta, Omicron BA1, BA2 and BA4/5 variants. Collectively, MAP-based DNA constructs encoding chimeric RBDs of SARS-CoV and SARS-CoV-2 variants, as represented by MAP-1002, are potential COVID-19 vaccine candidates worthy further translational study.

## Introduction

Effective vaccines against infection from the severe acute respiratory syndrome coronavirus 2 (SARS-CoV-2) are crucial weapons to control the ongoing pandemic coronavirus disease 2019 (COVID-19), which has caused more than 630 million infections with more than 6.5 million deaths worldwide since late 2019 (1). To date, more than 30 first-generation vaccines based on the ancestral (wild type, WT) strain of SARS-CoV-2 and several second-generation vaccines based on SARS-CoV-2 variants of concerns (VOCs) have been approved or authorized for emergency use, including inactivated virus vaccines, viral vector vaccines, subunit vaccines and nucleic acid vaccines encoding the viral spike (S) protein (2). Significantly decreased protective efficacies against emerging variants were observed in clinical trials and real-world evidence studies of first-generation COVID-19 vaccines (2–5). Waves of breakthrough infections of SARS-CoV-2 VOCs, including the ongoing Omicron BA.5 and BQ1.1 outbreaks around the globe, in previously vaccinated patients have been reported (5–11). It is thus necessary to develop novel vaccines able to provide broader-spectrum protection against newly emerging SARS-CoV-2 VOCs.

Among the four structural proteins in SARS-CoV-2, S protein is the main target for COVID-19 vaccines. It contains the receptor-binding domain (RBD) responsible for human ACE2 receptor binding and mediating virus entry (12,13). Neutralizing antibodies (NAbs) specific for RBD in S1 region of the S protein play critical roles in COVID-19 protection (14,15). Protein subunit vaccines based on recombinant WT SARS-CoV-2 tandem-repeat dimeric RBD or WT-Beta and Delta-Omicron BA1 chimeric RBD dimers induced NAb production and provided cross-protection against SARS-CoV-2 VOCs in mice and rhesus monkeys (16,17). Tan et al reported that BNT162b2 mRNA vaccine generated pan-Sarbecovirus NAbs in SARS-CoV survivors, suggesting that SARS-CoV-induced immunological memory cells could help production of broadly cross-reactive NAbs against SARS-CoV-2 variants (18). It is thus reasonable to speculate that combination of RBDs from pre-emergent SARS-CoV and SARS-CoV-2 variants could generate stronger immunogens able to further broaden the spectrum of cross-protection against SARS-related viruses.

DNA vaccines are considered an attractive alternative to conventional vaccines because they are relatively easy and inexpensive to produce, stable at room temperature, and able to stimulate potent cellular as well as humoral immunity (19,20). Several groups have explored the possibility to develop various DNA vaccines against COVID-19 with promising results. For example, COVID-eVax, an electroporated DNA vaccine candidate encoding the ancestral SARS-CoV-2 RBD, elicited protective responses in animal models (21). pGX9501, a WT SARS-CoV-2 full length (FL) S protein-encoding electroporated DNA vaccine candidate, was able to elicit NAbs as well as IFN-γ^+^-CD4 and CD8 T cells against WT SARS-CoV-2 as well as Delta variant in volunteers aged between 18 and 60 years (22). In 2021, ZyCoV-D, a S protein-encoding DNA vaccine delivered by a needleless injector, was authorized for emergency use against COVID-19 in India (23). However, naked DNA plasmids administered alone are relatively poor in transfection ability and consequently show low level of immunogenicity in vivo (19,20). Electroporation (EP)-, or needless injection-, assisted delivery can significantly improve DNA immunization results, but such methods often cause pain or discomfort to the vaccinees and require special expertise to operate the equipment. One possible solution to this problem is microneedle array patch (MAP) technology which utilizes microscopic projection arrays on a plaster to deliver a vaccine in the form of a patch placed on the skin (24,25). Due to its immune-rich milieu, the skin is a unique vaccination site evolutionarily primed to respond to challenges leading to strong adaptive humoral and cellular immunity (26–28). Evidence accumulated so far indicates that MAPs can intradermally deliver DNA vaccines for satisfactory immunization in an easy-to-use and painless fashion (30–31).

This study was designed to firstly construct DNA vaccine candidates encoding dimeric RBD chimera of SARS-CoV and SARS-CoV-2 variants and assess their immunogenicity, and also explore the possibility of developing MAP-based RBD-chimera DNA vaccines that can effectively induce cross-neutralizing Abs against antigen-matched and mismatched SARS-COV-2 VOCs. Our results suggest that combination of the RBD chimera approach, DNA vaccination and MAP technology opens a new avenue to overcome the shortcomings of the current vaccines and greatly augment cross-protection and global accessibility of vaccines in the fight against COVID-19 pandemic.

## Materials and Methods

### DNA vaccine construction

The synthesis of cDNA encoding heterodimeric fusion RBDs of SARS-CoV-1, prototype SARS-CoV-2 (2019-nCoV strain IVDC-HB-01/2019, GISAID: EPI_ISL_402119) and its variant B.1.351 (Beta), Omicron BA.1 and BA.5 was performed by GenScript, Nanjing, China. Optimization analysis of the cDNA sequences was performed using an in-house analytic tool, taking into accounts codon usage bias, GC content, mRNA secondary structure, cryptic splicing sites, premature poly(A) sites, internal *chi* sites and ribosomal binding sites, negative CpG islands, RNA instability motif (ARE), repeat sequences (direct repeat, reverse repeat, and dyad repeat), and restriction sites that may interfere with cloning. The resulting synthesized and optimized cDNA, together with a secretion leader peptide-encoding sequence, was cloned into expression vector pVAX1. Plasmid pWT, a pVAX1-based COVID-19 vaccine candidate encoding FL S protein of prototype SARS-CoV-2, was as previously described (32). A pVAX1-based expression plasmid encoding full length (FL) firefly luciferase (pVAX1-Luc) was similarly constructed. Restriction enzyme analysis and DNA sequencing was performed to confirm the accuracy of construction. Plasmids were transformed into *E. Coli* strain HB101. Single colonies were undergone expansion in one-liter flasks for culturing in LB broth. Plasmids were extracted, purified by MaxPure Plasmid EF Giga Kit (Magen, China), and dissolved in distilled water at 1 mg/mL final concentration. Purity of the plasmids was measured by an agarose gel electrophoresis and a UV detector at a range of 1.8-2.0 OD260 nm/280 nm. Endotoxin contamination in plasmid samples was below 30 EU/mg by the LAL test.

### Fabrication of MAPs laden with DNA vaccine

MAPs used in this study were prepared by a two-step micro-molding process. The vaccine formulation consisting of concentrated DNA plasmid (adjusted to obtain a final dose of 20 μg per patch), water-soluble and biocompatible materials including polyvinyl alcohol (PVA), hydroxylethyl cellulose (HEC), polyvinyl pyrrolidone (PVP), sucrose and supplemental salts in 100 mM trisodium citrate buffer pH 7.4 was cast onto a PDMS mold (486 MNs per array; each cone-shaped MN measuring 450 μm in length and 160 μm in width at the base). Vacuum was applied to ensure that the formulation filled the entire MN cavity and the formulation was allowed to air dry at room temperature overnight. Then the backing formulation consisting of PVA, PVP and sucrose was cast onto the mold under vacuum and subsequently dried at room temperature for 4 h before demolding the MN patch, which was further mounted onto a 1.5 cm^2^ paper backing. The resulting patches had a moisture content less than 2% and were stored in sealed individual aluminum pouches with desiccant at +4°C or +25°C until use. The strength of MNs was checked before and after short-term insertion in a ten-layer parafilm pack for penetration effectiveness, bending and brittleness under the microscope.

### Western blot

HEK293 cells that had been pre-plated in a 6-well plate were transiently transfected with 2.5 μg DNA vaccine plasmids with Hieff TransTM Liposomal Transfection Reagent (YEASEN, Shanghai, China). Two days later, the cells were pelleted and lysed in immunoprecipitation assay buffer. Cell lysates and supernatants were separated by SDS-PAGE and transferred to PVDF membranes. Immunoblotting was performed by using rabbit anti-RBD^WT^ primary Ab (Bioworld, Nanjing, China) diluted 1:1,000 in 5% milk-0.05% PBS-Tween 20, and horseradish peroxidase (HRP)-labeled goat anti-rabbit IgG secondary Ab (BD Biosciences, San Diego, CA, USA). Chemiluminescence detection was performed with the ECL Prime Western Blotting System and acquired by the ChemiDoc Imaging System (Bio-Rad).

### Bioluminescence Imaging

BALB/c and C57BL/6 mice (5 mice/group) were anesthetized with 97% oxygen and 3% isoflurane (Isoba, MSD Animal Health, Walton, UK) and then administered with MAP-Luc patches on shaved skin surface on dorsal sides for 15 min. Fifteen min after i.p. injection of a 15 mg/mL luciferin solution (PerkinElmer) at 10 μL/g body weight, the mice were subjected to bioluminescence imaging using IVIS Spectrum under gas anesthesia. Luciferase expression level was then quantified using the Living Image software in a fixed region of interest (ROI) in terms of photone/sec/cm2/sr.

### qRT-PCR

SARS-CoV-2 RBD-specific quantitative reverse transcription-PCR (qRT-PCR) assays were performed by using a FastKing One Step Probe RT-qPCR kit (Tiangen Biotech, China) on a CFX96 Touch real-time PCR detection system (Bio-Rad, USA).

### Animal immunization

Female BALB/c and C57BL/6 mice (6-8 weeks of age) were purchased from Shanghai Slac Laboratory Animal Co., Ltd. and maintained under SPF conditions at the animal facilities of Advaccine Biologics (Suzhou) Co. New Zeeland white rabbits, purchased from Shanghai Somglian Experimental Animal Company, were housed in the Grade I animal facilities of Advaccine Biologics (Suzhou) Co. All animal experiments were performed in compliance with the recommendations in the Guide for the Care and Use of Laboratory Animals of the Ministry of Science and Technology Ethics Committee and approved (document No. 2021070102) by on the Ethics Committees of the company.

EP-assisted DNA immunization was performed in mouse and rabbit quadriceps injected with 20 μg (for mice) or 0.5 mg (for rabbits) DNA (1 mg/mL in 30 μl SSC), followed by EP using Inovio CELLECTRA^®^2000 and needle electrode (Inovio, San Diego, CA, USA) with two sets of pulses with 0.2 Amp constant current. All IM + EP-delivered vaccines were primed on day 0 and boosted on day 14 unless otherwise indicated. Blood samples were collected on days 0, 14, 21 and/or 28.

To administer MAP-based DNA vaccines in mice and rabbits, patch-sized skins on dorsal sides were shaved and treated with hair removal cream one day earlier. Apply and slightly press the patch with thumb pressure on the shaved skin surface of the animals under anesthesia, allow to stay for 15 minutes and then peel off the used MAP. MAPs fabricated in this study did not cause skin allergy or physical damage, and within 6 days fur of the shaved skins returned to normal.

### Enzyme-linked immunosorbent assay

Antibody titration was performed on sera obtained by retro-orbital bleeding from mice or venous bleeding from the ears of rabbits. The ELISA plates were functionalized by coating with the recombinant RBD proteins (SinoBiological, Beijing, China) at 1 μg/mL and incubated 18 h at 4°C and subsequently blocked with 3% BSA-0.05% Tween 20-PBS (PBST) for 1 h at room temperature. Serial diluted serum samples were then added in triplicate wells, and the plates were incubated for 1 h at room temperature. After a double wash with PBST, horseradish peroxidase (HRP)-conjugated Ab against murine (Abcam, ab6789, 1/2000 diluted), or rabbit (GenScript, A00098, 1:2,000 diluted) IgG was added and then developed with 3,3’,5,5’ -tetramethylbenzidine (TMB) substrate (Coolaber, CN). The reaction was stopped with 2 M of H_2_SO_4_, and the absorbance measured at 450 nm and reference 620 nm using a microplate reader (TECAN, CH).

### Neutralization antibody detection

The pseudo-virus microneutralization assay was performed to measure neutralizing antibody levels against prototype SARS-CoV-2 and its variants. Pseudovirus stocks of prototype SARS-CoV-2, variants and B.1.35, P.1, B.1.617.2 were purchased from Gobond Testing Technology, which were aliquoted for storage at −80 °C. hACE2 stable expressing HEK293T cells (prepared in our lab) were used as target cells plated at 10,000 cells/well. SARS-CoV-2 pseudo-viruses were incubated with heat-inactivated (56°C for 30 min) and 1/3 serial diluted mouse sera for 90 min at room temperature; then, the sera-pseudotype-virus mixtures were added to hACE2-HEK293T cells and allowed to incubate in a standard incubator 37% humidity, 5% CO_2_ for 72 h. The cells were then lysed using Bright-Glo™ Luciferase Assay (Promega Corporation, Madison, WI, USA), and RLU was measured using an automated luminometer. Fifty percent pseudovirus neutralization titer (pVNT50) was determined by fitting nonlinear regression curves using GraphPad Prism and calculating the reciprocal of the serum dilution required for 50% neutralization of infection. These assays have been performed in a BSL-2 facility of Advaccine. Pseudovirus neutralization experiments using Vero cells were contracted to Gobond Testing Technology, Beijing, China.

### ELISpots

Spleens and draining lymph nodes (LNs) from immunized mice were collected and used to prepare single cell suspension in RPMI-1640 medium supplemented with 10% FBS and penicillin/streptomycin. ELISpot was performed using mouse IFN-γ and IL-4 ELISpot PLUS kits (MABTECH, Cincinnati, OH, USA) according to the manufacturer’s protocol. Briefly, 5×10^5^ freshly prepared mouse splenocytes, or LN cells, were plated into each well and stimulated for 20 h with pooled overlapping 15-mer peptides (10 μg/ml) covering respective RBDs at 37°C in a 5% CO_2_ incubator. PMA/Iono was used for positive controls. The plates were processed in turn with a biotinylated detection antibody. Spots were scanned and quantified using AID ImmunoSpot reader (AID, GER). IFN-γ- and IL-4-spot forming units were calculated and expressed as SFUs per million cells.

### Intracellular cytokine stain (ICS)

Freshly isolated mouse splenocytes or LN cells were stimulated with an overlapping peptide pool of RBD^WT^ (10 μg/mL) in the presence of Brefeldin A (Invitrogen, USA) for 5 h at 37 °C, 5% CO_2_. The cells were harvested and stained with anti-CD3, anti-CD4 and anti-CD8α surface markers, and subsequently fixed and permeabilized in permeabilizing buffer (eBiosciences, USA) and stained with fluorescence-conjugated anti-IFN-γ, anti-TNF-α and anti-IL-2 and anti-IL-4 antibodies. All the fluorescence-labeled Abs were from BioLegend, and the stained lymphocytes were analyzed on Attune NxT Flow Cytometer (ThermoFisher, USA). The data were analyzed by Attune™NxT software (ThermoFisher, USA).

### Statistics

Statistical analyses were performed with GraphPad Prism software version 9 (GraphPad). Error bars indicate the standard error of the mean (SEM). We used Mann-Whitney t-tests to compare two groups with non-normally distributed continuous variables and two-way ANOVA followed by Sidak’s multiple comparisons tests to analyze experiments with multiple groups and two independent variables. Significance is indicated as follows: *p < 0.05; **p < 0.01. Comparisons are not statistically significant unless indicated.

## Results

### Preparation and immunogenicity evaluation of DNA constructs encoding dimeric RBD chimera of SARS-CoV and SARS-CoV-2 variants

Three pVAX1-based COVID-19 vaccine candidates encoding heterodimeric fusion RBDs between SARS-CoV-1 (GenBank accession no: AY278488.2) and SARS-CoV-2 variant Beta (EPI_ISL_860630, GISAID) or Omicron BA.1 (EPI_ISL_6640917, GISAID), namely pADV131 (SARS-Beta), pAD1002 (SARS-Omicron) and pAD1003 (Beta-Omicron), were constructed (**Fig. 1A**). RNA- and codon-optimization was performed to increase the expression efficiency of the DNA constructs in mammalian cells. To promote protein secretion, we introduced a unique secretion leader sequence in the fusion RBD constructs. Expression of the resultant plasmids in transfected HEK293T cells was confirmed by qPCR and Western blotting (**Figs. 1B & 1C**). Secreted recombinant RBD proteins were readily detectable in culture supernatant of the transfectant cells using ELISAs (**Figs. 1D–1F**). For immunogenicity evaluation, groups of BALB/c mice were intramuscularly (IM) administered with 2 doses (20 μg/dose, with fortnight interval) of the plasmids, followed by ELISA monitoring of serum IgG against recombinant RBD of WT SARS-CoV-2 (RBD^WT^). As shown in **Fig. 1G**, i.m. inoculated plasmids pAD1002 and pADV131 were able to induce reasonably strong IgG responses in mice. By contrast, however, i.m. injected pAD1003 was essentially non-immunogenic. This is similar to the FL S protein-encoding plasmid pWT, which is also a poor immunogen when IM administered alone and requires EP assistance to trigger decent humoral responses in vivo (**Fig. 1H**). Given that pAD1002 and pADV131 differ from pAD1003 and pWT in possessing SARS-CoV RBD (RBD^SARS^)-encoding sequence, these results argue for potent immunogenicity-boosting effect of RBD^SARS^ in vivo.

**Fig. 1.**
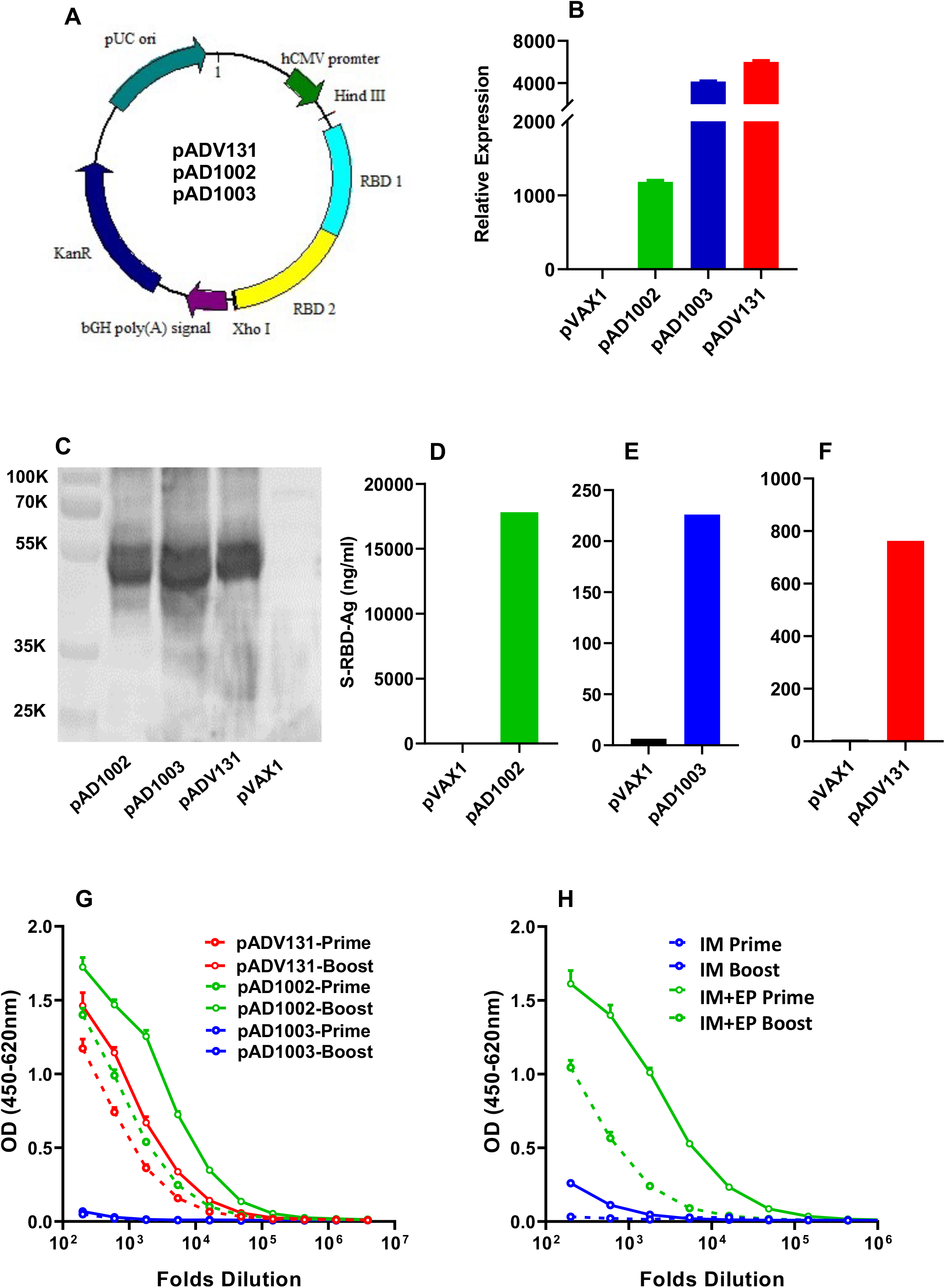
Construction and immunogenicity evaluation of COVID-19 vaccine candidates. (**A**) Schematic diagram showing the structure of pVAX1-based vaccine candidates encoding heterodimeric fusion RBDs of SARS, Beta and Omicron BA1 with a secretion leader sequence. Gene fragments were cloned into the BamH1 and XhoI sites of pVAX1 vector under human CMV promoter control. HEK293T cells, transiently transfected with pVAX1, pAD1002, pAD1003 or pADV131 24 h earlier, were subjected to (**B**) qPCR detection of RBD mRNA transcripts using GPDH as internal control, and (**C**) SDS-PAGE gel electrophoresis followed by Western blotting using anti-RBD^WT^ Abs for detection. (**D-F**) Secreted recombinant RBD chimera in culture supernatant of the transfectant HEK293T cells were quantitated using RBD^WT^-based ELISAs. (**G**) Serum samples, collected from BALB/c mice after primary and secondary i.m. immunizations with plasmid pAD1002, pAD1003 or pADV131 (20 μg/dose) were titrated against recombinant RBD^WT^ in ELISAs. (**H**) Sera from BALB/c mice 14 days after secondary IM or IM + EP inoculation of plasmid pWT were analyzed in RBD^WT^-based ELISA. Data represent mean ± SD (n=3 biologically independent samples).

### Fabrication and characterization of MAPs for intradermal DNA delivery

We next sought to develop novel COVID-19 vaccine preparations by combining the RBD chimera DNA with MAP technology, which may bypass the need of EP-assisted delivery of DNA vaccines for satisfactory immunization results. The two-step micro-molding procedure to fabricate Advaccine MAP (MAP^Adv^) is shown in **Fig. 2A**, which has consistently given sharp and robust MN structures capable of penetrating stratum corneum with thumb pressure. The resulting round-shaped skin patch is arranged in a 486 MN array covering an area of 1.5 cm^2^ (**Fig. 2B**). Microscopic examination confirmed that arrayed MNs on the resultant patch are 550 μm in height, including 450 μm cone-shaped needle and 160 μm base. DNA plasmids are entrapped in the top third region of the dissolvable MNs, the width of pinpoint is less than 10 μm and the tapered base 160 μm (**Figs. 2C & 2D**). Once inside the epidermis, MN tips readily dissolve to release the DNA load within minutes. Individually bagged with desiccant, MAP^Adv^ patches laden with 20 μg plasmid DNA of pAD1002, pAD1003 or pADV131, namely MAP-1002, MAP-1003 and MAP-131, respectively, are structurally and functionally stable in room temperature and do not require refrigeration for long term storage. After 30 days of storage at 25°C, for example, over 98.5% plasmid DNA recovered from MAP-1002 remained in a supercoiled form (**Figs. 2E & 2F**), and the MAP retained potency for intradermal immunization (see below).

**Fig. 2.**
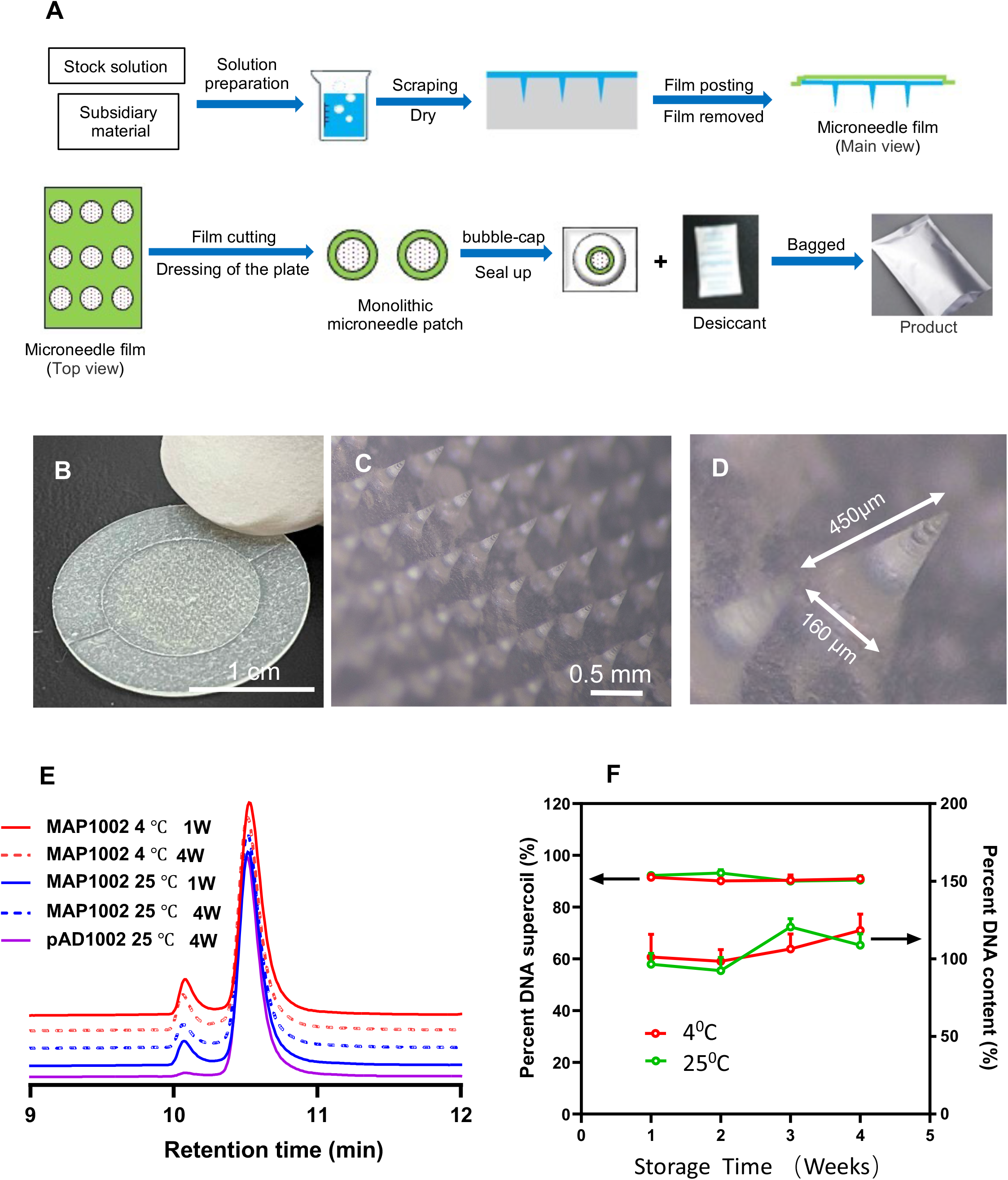
Fabrication of MAP^Adv^ for intradermal DNA delivery. (**A**) Schematic diagram showing the fabrication procedure for MAP^Adv^. (**B**) Photograph of a MAP^Adv^ skin patch, which is of 1.2 cm in diameter, with an array of 486 MNs loaded with 20 μg plasmid DNA. (**C**) Microphotograph showing the structure of arrayed MNs in MAP^Adv^. The MNs in MAP^Adv^ patch are 450 μm in height, including 320 μm cone-shaped needles and tapered base 130 μm. For stability analysis, samples of MAP-1002 patches, maintained at 4° or 25° for 1-4 weeks, were emersed in 2 ml saline for 1 h to allow complete dissolution of MNs, then the solution was subjected to HPLC profiling (**D**), and quantitation of DNA (expressed as ratio of recovered/theoretical DNA) and percent plasmid DNA in supercoil form (**E**).

### Intradermal expression of gene delivered using MAP^Adv^

Luciferase (Luc) activity in living animals can be visualized by bioluminescence imaging in the presence of D-luciferin. To assess the expression efficiency of MAP^ADV^-delivered genes in vivo, we constructed a pVAX1-based expression vector encoding FL firefly Luc (pVAX1-Luc) and then fabricated MAP-Luc patches (20 μg DNA/patch). BALB/c mice were treated with the resultant MAP-Luc patches on dorsal sides, followed by bioluminescence imaging 24 h thereafter. Strong bioluminescence signals were observed at MAP-Luc application sites (**Fig. 3A**). Forty-eight h after MAP-Luc treatment, mice were sacrificed for skins, skinned bodies, spleens and draining LNs (dLNs) which were immediately subjected to bioluminescence imaging. Bioluminescence signals were detected only on sites of patch application in the skins, not the skinned bodies, spleens or dLNs (**Fig. 3B**).

**Fig. 3.**
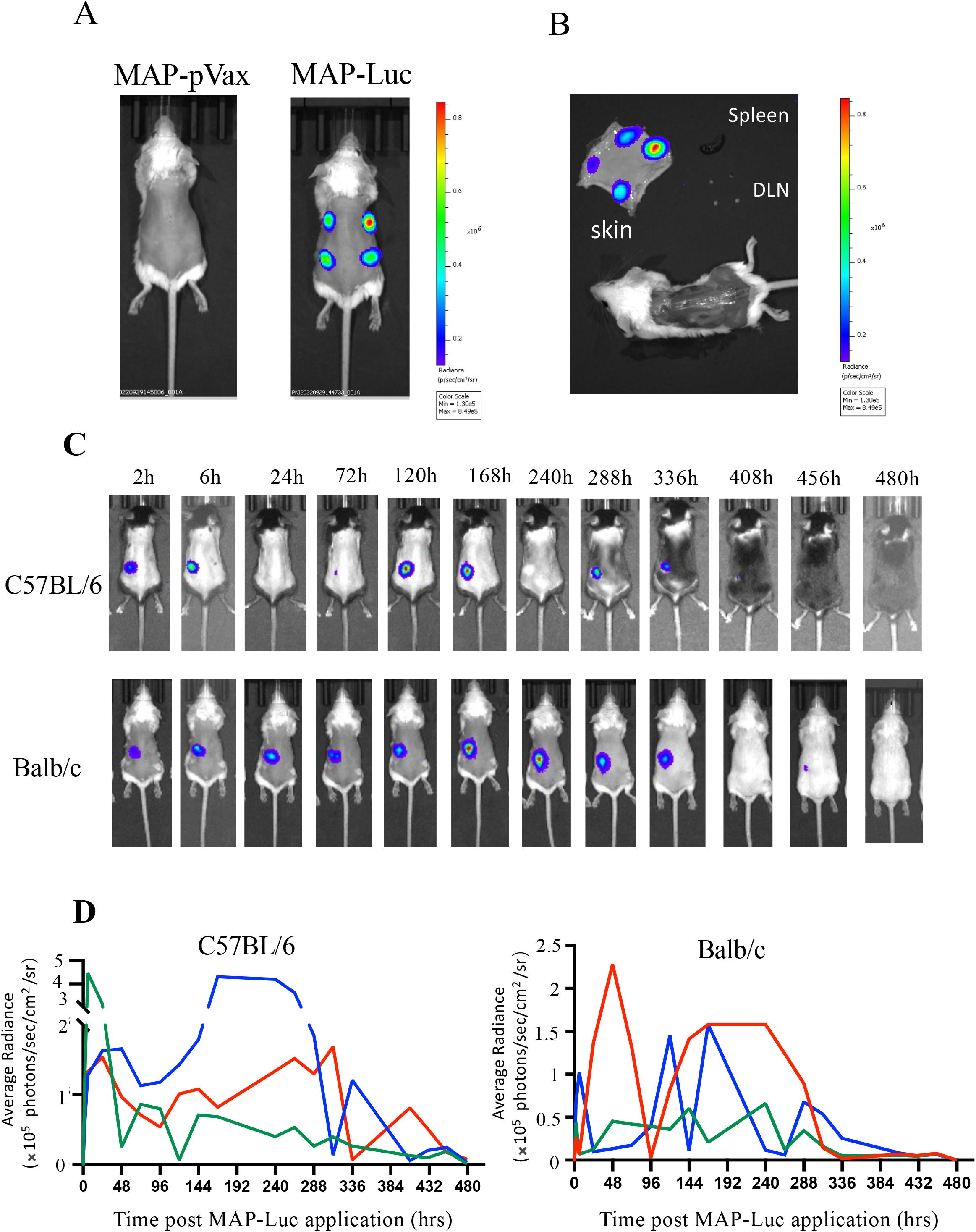
In vivo visualization of gene expression in mouse skin after MAP^Adv^-mediated pVAX1-Luc delivery. (**A**) Bioluminescent images of living BALB/c mice administered with 4 MAP-Luc, or MAP-pVAX1, patches on dorsal sides 24 h previously. (**B**) Freshly prepared dorsal skin, skinned body, dLNs and spleen from a BALB/c mouse treated with 4 MAP-Luc patches 48 h earlier were subjected to bioluminescent imaging. (**C**) C57BL/6 and BALB/c mice (n=3) were administered with MAP-Luc patches (1 patch each mouse, on dorsal side, for 15 min), followed by bioluminescent imaging at different timepoints up to 480 h. Images from one representative mouse of each group are shown. (**D**) Bioluminescence intensity at sites of MAP-Luc application for each mouse was quantitated and recorded. The above is representative of 3 independent experiments.

To study the expression kinetics of MAP-delivered DNA in vivo, groups of BALB/c and C57BL/6 mice were treated with MAP-Luc patches and then monitored for Luc gene expression (biofluorescence) at different timepoints. As illustrated in **Fig. 3C**, strong bioluminescence signals were readily detectable at the patch application sites as early as 2 h post MAP-Luc administration in both strains of mice, lasting for 15 days or more. Considerable individual variation in intradermal gene expression profile after MAP-Luc administration was observed, particularly in C57BL/6 mice. In general, all MAP-Luc-treated mice expressed Luc gene in waves of hrs or days before the fluorescence signals eventually dropped below detection (**Fig. 3D**). The strong and durable intradermal expression of MAP^Adv^-delivered gene in BALB/c and C57BL/c mice provides solid basis for potent immunological responses against the target antigens triggered by MAP-DNA vaccination in the hosts.

### Immunogenicity-boosting effect of MAP^Adv^ in DNA immunization

EP has so far been the most effective, albeit rather uncomfortable, method to enhance DNA vaccine immunogenicity in vivo. It was of interest to know if MAP^Adv^ can be as efficient as IM + EP in enhancing DNA vaccination efficiency in vivo. BALB/c mice were administered with two doses of MAP-1002, MAP-1003, or MAP-131 (1 patch/dose/mouse, with a fortnight interval). For controls, two doses of corresponding naked plasmids (20 μg/dose) were IM injected with, or without, EP. Results (serum IgG titers) of RBD-DNA immunization mediated by MAP^Adv^, IM or IM + EP delivery are compared in **Figs. 4A–4C**. After boost, RBD^WT^-binding IgG titers of the MAP^Adv^ and IM + EP groups were nearly identical. MAP (and IM + EP) delivery of plasmid pAD1002 and pADV131 led to 5 folds higher anti-RBD IgG titers compared to IM immunization alone. The most significant MAP enhancement in antibody response to DNA immunization was seen in MAP-1003, which completely overcome the very poor immunogenicity of IM-inoculated plasmid pAD1003 (**Fig. 4B**). RBD-specific serological IgG titers in both the MAP^Adv^ and IM + EP groups maintained at high levels for 70-180 days post immunization (**Figs. 4D–4F**). A dose-response curve of MAP-mediated DNA vaccination was obtained by plotting RBD^WT^-binding IgG titers against decreasing DNA doses delivered by a full-, half-or quarter-sized MAP-131 (**Fig. 4G**). MAP-1002 was also employed to immunize C57BL/6 mice (1 patch/dose/mouse) and New Zeeland white rabbits (10 patches/dose/rabbit), with MAP-pVAX1 and pAD1002/IM + EP as controls. Strong and lasting IgG responses to MAP-1002 and pAD1002/IM + EP immunizations were also observed (**Figs. 4H & 4I**). Therefore, MAP^Adv^ represents an effective alternative to IM + EP for enhancing the results of DNA vaccination in experimental animals of different genetic backgrounds.

**Fig. 4.**
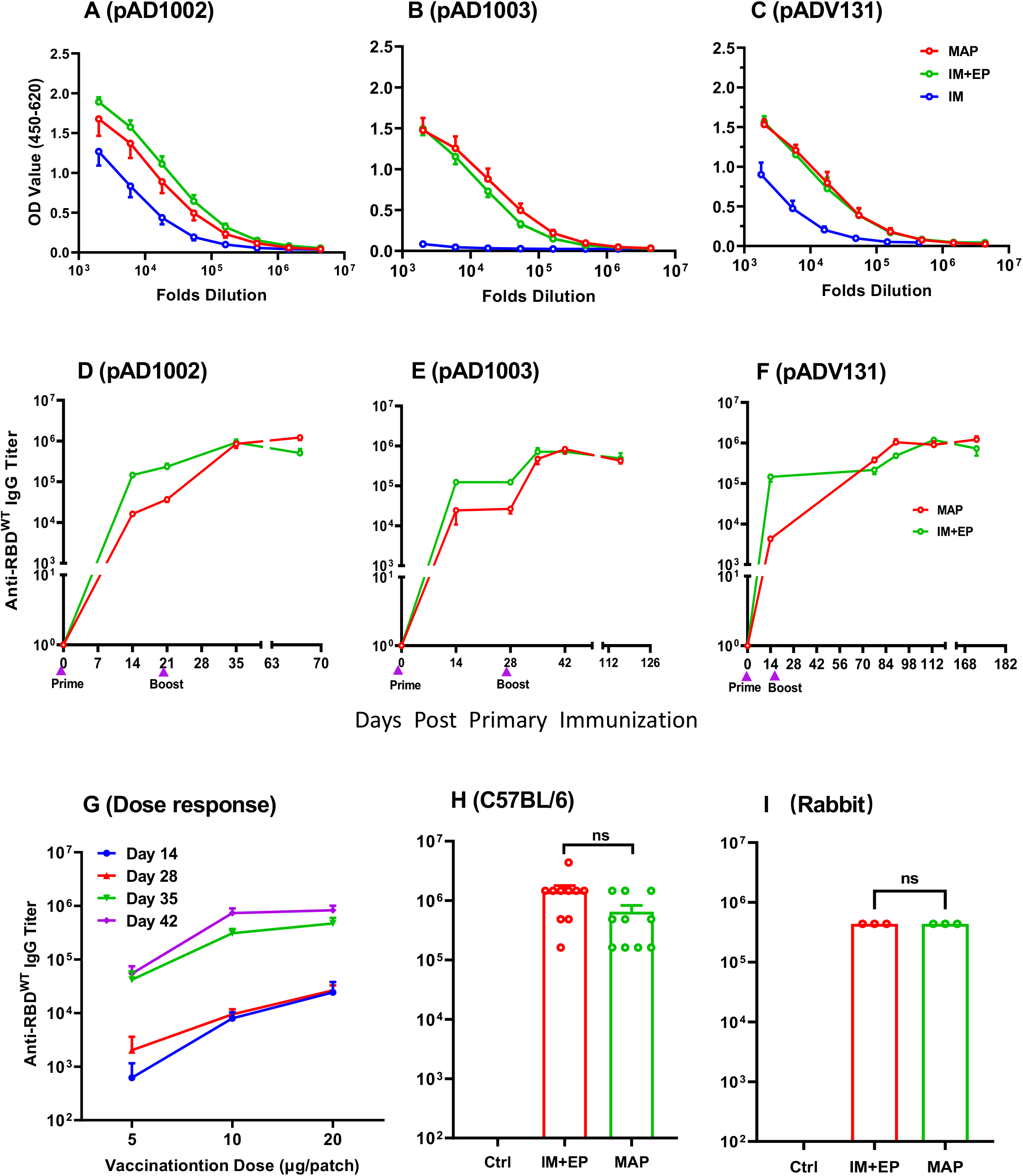
Immunogenicity-boosting effect of MAP^Adv^ in DNA immunization. Groups of BALB/c mice were given two doses (20 μg DNA/dose, with fortnight interval) of plasmid pADV131 (**A**), pAD1003 (**B**), or pAD1002 (**C**) via MAP, IM or IM + EP delivery. Serum samples, collected 14 days after boost, were titrated against recombinant RBD^WT^ in ELISAs. Serum IgG titers in BALB/c mice after MAP- or IM + EP-mediated immunization with plasmid pADV131 (**D**), pAD1002 (**E**), or pAD1003 (**F**) were monitored using RBD^WT^-based ELISAs for up to 120 days after immunization. (**G**) Serum samples, collected from BALB/c mice (n = 10) on days 14, 28, 35 and 42 after primary (Day 0) and secondary (Day 14) immunization with decreasing DNA doses (20, 10, 5 μg/dose) delivered by a full-, half- or quartersized MAP-131, respectively, were titrated against recombinant RBD^WT^ in ELISAs. Dose response curve was obtained by plotting end-point dilution serum IgG titers against delivered DNA doses. (**H**) Serum IgG titers in C57BL/6 mice (n=5) and (**I**) New Zeeland white rabbits (n=3) administered with two doses of MAP-1002 (1 patch/dose/mouse, 10 patches/dose/rabbit), or pAD1002/IM + EP (20 μg/dose/mouse, 200 μg/dose/rabbit), or pVAX1/IM + EP as control. Data represent mean ± SEM. Unpaired t-test was used to determine significance within **F** and **G** (*P< 0.05, **P< 0.01, ***P< 0.001, ****P< 0.0001).

### Virus-specific CTL responses elicited by MAP-1002 in mice

Compared to inactivated virus or subunit viral protein vaccines, nucleic acid vaccines are particularly powerful in generating MHC I-restricted CD8^+^ cytotoxic T lymphocytes (CTL) known to play pivotal roles in protection against viral infections in vivo (20). SARS-CoV-2 virus-specific CTL responses have been found to be associated with milder situations in acute and convalescent COVID-19 patients (33). To evaluate the ability of MAP-DNA vaccine candidates to induce CTLs in vivo, C57BL/6 mice were administered with 2 doses of MAP-1002, inactivated WT SARS-CoV-2 vaccine, or pVAX1/IM + EP as sham control, followed by ELISpot detection of RBD-responding IFN-γ^+^ and IL-4^+^ cells in spleens and dLNs. Interestingly, RBD-responsive IFN-γ^+^ cells were found in dLNs and, to a lesser extent, spleens of mice immunized with MAP-1002, but not pVAX1 or inactivated virus vaccine (**Fig. 5A**). On other hand, IL-4^+^ cells elicited by MAP-1002 were found mostly in spleens rather than dLNs, while inactivated virus vaccine generated IL-4^+^ cells detectable in both spleens and LNs (**Fig. 5B**). This is in line with the concept that inactivated virus vaccines mainly induce humoral responses while MAP-mediated DNA immunization elicits balanced cellular (CTL) and humoral immunity. EP-assisted DNA immunization is also well known for ability to trigger strong T cell immunity in vivo (20). Interestingly, pAD1002/IM + EP immunization of BALB/c mice generated RBD-responsive IFN-γ^+^ cells almost exclusively present in the spleens rather than LNs (**supplemental Fig. S1**), which is contrasting the “dLN-favoring” distribution pattern of RBD-specific T cells induced by MAP-1002 in mice. Intracellular cytokine staining (ICS) results confirmed that RBD-specific IL-2-, IFN-γ- and TNF-a-expressing CD8^+^ T cells were clearly identifiable in dLNs from BALB/c mice 14, 21 and 35 days post MAP-1002 immunization (**Figs. 5C–5E**). By Day 35, percentage of CD8^+^ T cells bearing CX3CR1, a surface marker for effector memory CD8^+^ T cells (TEM), in dLNs of MAP-1002-immunized mice were significantly higher compared to that of the pAD1002/IM or unimmunized control groups (**Figs. 5F & 5G**).

**Fig. 5.**
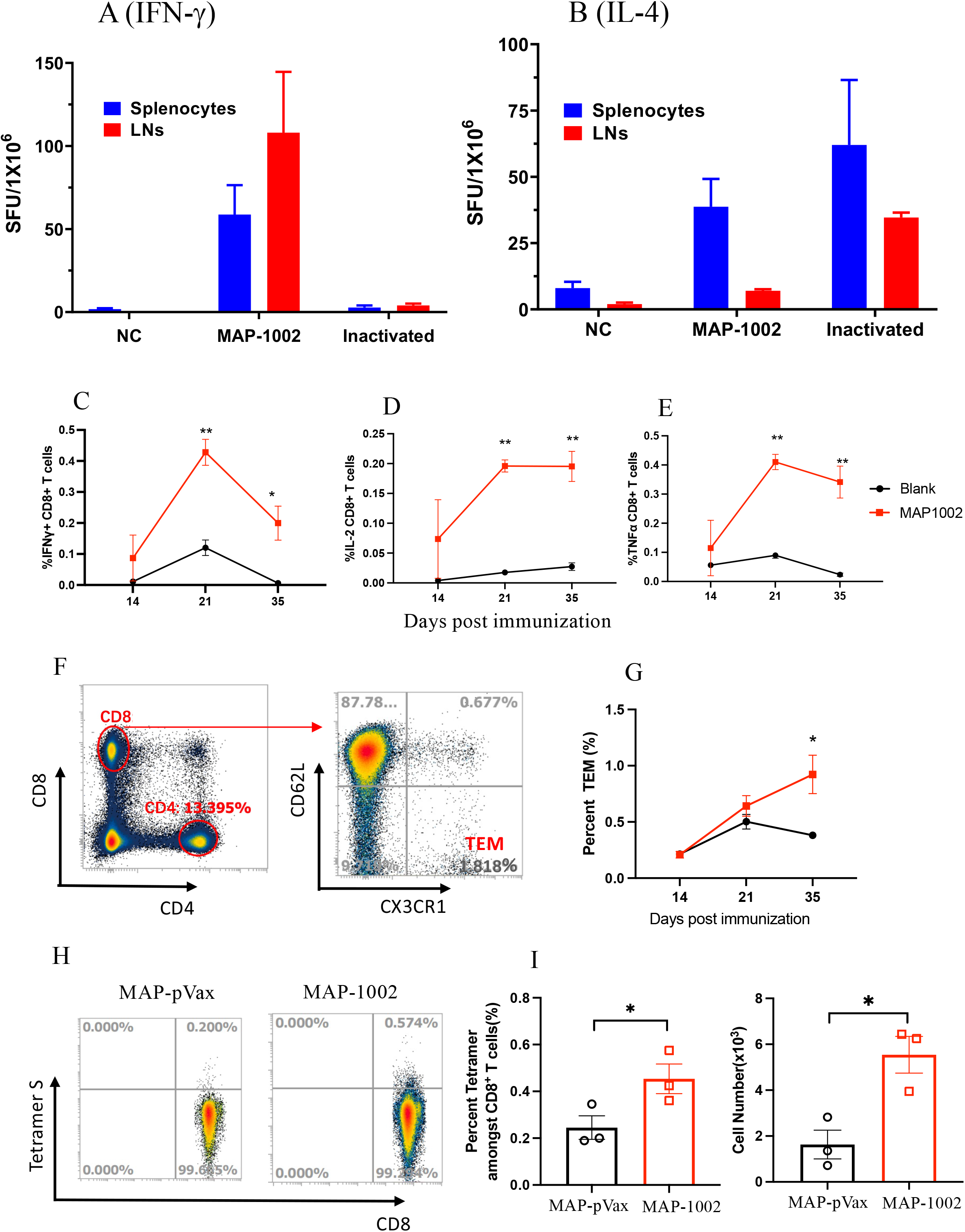
SARS-CoV-2-specific T cell responses in MAP-DNA-vaccinated mice. BALB/c mice were immunized twice with, or without (Blank), MAP-1002 or inactivated virus vaccine (Inact/IM) and then sacrificed for spleens and dLNs 14 days after boost. ELISpot analyses of (**A**) IFN-γ and (**B**) IL-4 spot-forming cells (SFC) in splenocytes and dLN cells after re-stimulation with pooled 14-mer overlapping peptides spanning the RBD^WT^ sequence were performed. (**C-E**) CD8^+^ T cells amongst dLN cells of BALB/c mice 14, 21 and 35 days post MAP-1002 immunization were assayed for IFN-γ, IL-2 and TNF-a expression by CIS and FACS analysis after re-stimulation with the RBD^WT^ peptide pool. LN cells from the unimmunized (blank) mice were included as control. (**E**) Percentage of CD8^+^ TEM in LN cells of BALB/c mice 14, 21 and 35 days after MAP-1002 administration, as revealed by flow cytometric analysis results, are compared with that of unimmunized mice (Blank). (**H-J**) Draining LN cells from C57BL/6 mice 14 days after MAP-1002 immunization were stained with PE-labeled “S Tetramer”for identification of CD8^+^ T cells expressing TCR specific for S peptide/ H-2D^b^ complex by flow cytometry. Data are means ± SEM. Absolute number of S Tetramer-binding cells and their percentage amongst CD8^+^ cells are shown in the histograms. P values were analyzed with two-tailed Mann-Whitney test (ns, P > 0.05; *P < 0.05; **P < 0.01).

Several mouse CTL epitopes have been identified in SARS-CoV-2 RBD, including a H-2D^b^-restricted “S”epitope (amino acid residues 366-374, SVLYNSASF) (34). We next employed the *S* tetramer (PE-labeled H-2D^b^ tetramer harboring the “S” peptide), in flow cytometric analysis, to capture CD8^+^ T lymphocytes bearing TCRs capable of recognizing the *S* epitope amongst dLN cells from MAP-1002-primed C57BL/6 mice. As illustrated in **Figs. 5H & 5I**, the LN-residing S epitope-specific CTLs expanded significantly as result of MAP-1002 immunization in C57BL/6 mice. Collectively, MAP-1002 was able to generate RBD-specific CD8^+^ CTL responses in mice, and the resulting T lymphocytes homed mainly in dLNs rather than spleens of the responder animals.

### Broadly cross-binding property of serological IgG induced by MAP-based DNA vaccine candidates encoding heterodimeric RBDs

The heterodimeric RBD approach was designed to broaden the spectrum of immune protection against antigen-matched and antigen-mismatched SARS-CoV-2 VOCs. To evaluate the cross-reactivity of MAP^Adv^-DNA-generated mouse IgG antibodies, ELISAs were carried out against a panel of 6 recombinant RBD preparations representing RBD^WT^, RBD^Beta^, RBD^Delta^, RBD^BA1^, RBD^BA4/5^ and RBD^SARS^. Antisera from pWT-immunized mice were included as control. The results are presented in **supplemental Fig. S2** and RBD-binding titer and spectra of serological IgG from the 4 immunization groups are further compared in **Table 1**. Here RBD^WT^ and RBD^Delta^ can be regarded as “antigen-mismatched” viral antigens, as they are not encoded by any of the 3 MAP-based DNA vaccine candidates. Interestingly, both these two recombinant RBDs were strongly bound by serum IgG from the MAP-1002, MAP-1003 and MAP-131 groups (average endpoint dilution titers 486,000-1,458,000). Since late 2021, Omicron subvariants BA.4 and BA.5 have been circulating globally and gradually substituted its predecessors BA.1 and BA.2. Mutations in the BA.4/5 spike protein led to resistance for humoral immune responses induced by early SARS-CoV-2 infection and vaccination (5–11). Interestingly, both recombinant RBD^BA1^ and RBD^BA4/5^ were strongly bound by serological IgG of MAP-DNA-vaccinated mice (average endpoint dilution titers 378,000-486,000). Meanwhile, genetic distance between the immunizing DNA and coating RBD antigens did impact the cross-binding ELISA results. For example, RBD^Beta^-binding titers of the MAP-1003 and MAP-131 (both RBD^Beta^-encoding) antisera were over 3 times of that of the MAP-1002 (non-RBD^Beta^-encoding) antisera. RBD^SARS^-binding titer of the MAP-1003 (non-RBD^SARS^-encoding) antisera was less than 1% of the MAP-1002 and MAP-131 (both RBD^SARS^-encoding) antisera. It is also of importance to note that pWT-generated antisera did not bind RBD^BA1^, RBD^BA4/5^, or RBD^SARS^, and their binding titers to RBD^WT^ and RBD^Delta^ were some 2-10 folds lower compared to that of the MAP-DNA groups. These data support the idea that heterodimeric RBD approach can improve immunogenicity and help to expand the cross-reaction spectra of COVID-19 vaccines.

**Table 1.** Comparison of RBD cross-binding titers of serum IgG in BALB/c mice following immunization with MAP-based DNA vaccines or pWT/IM + EP^a^)

| Coating Ags<sup>b)</sup> | MAP-1002<sup>c)</sup> | MAP-1003<sup>c)</sup> | MAP-131<sup>c)</sup> | pWT/IM+EP<sup>c)</sup> |
| --- | --- | --- | --- | --- |
| RBD<sup>WT</sup> | 100 | 160.30±13.09 | 344.60±29.53 | 40.97±6.15 |
| RBD<sup>Beta</sup> | 100 | 313.27±9.69 | 329.02±19.31 | 10.53±0.71 |
| RBD<sup>Delta</sup> | 100 | 73.89±4.13 | 88.97±5.40 | <10 |
| RBD<sup>BA1</sup> | 100 | 255.45±25.33 | 74.38±9.12 | <5 |
| RBD<sup>BA4/5</sup> | 100 | 128.67±5.32 | 174.75±21.96 | <5 |
| RBD<sup>SARS</sup> | 100 | <5 | 76.98±4.82 | <1 |

### Pseudo-virus cross-neutralization by antisera from mice immunized with MAP-based DNA vaccines encoding RBD chimera

Generation of NAbs is known to be crucial for protecting people from virus infection. NAb levels are highly predictive of immune protection from symptomatic SARS-CoV-2 infection in humans (14,15). It was therefore of importance to ascertain if the high titer RBD-binding Abs induced by MAP-DNA vaccination in model animals positively correlate to virus neutralization capability. Firstly, we employed pseudo-viruses displaying recombinant S proteins of prototype SARS-CoV-2, or Omicron subvariant BA.1, to mimic infection of ACE2-expressing HEK293T cells. For WT SARS-CoV-2 pseudo-virus blocking, antisera from mice immunized with MAP-1003, MAP-1002, or MAP-131 were all effective (**Fig. 6A**), supporting the notion that heterodimer RBD approach can help to generate cross-protective immunity in vivo. For Omicron BA.1 pseudo-virus blocking, however, MAP-131 antisera were significantly poorer compared to MAP-1003 and MAP-1002 antisera (**Fig. 6B**), which is underlined by the fact that construct pADV131 does not encode Omicron RBD and that RBD^BA1^-binding titer of the MAP-131 antisera was lower than that of the MAP-1002 and MAP-1003 antisera (**Table 1**).

**Fig. 6.**
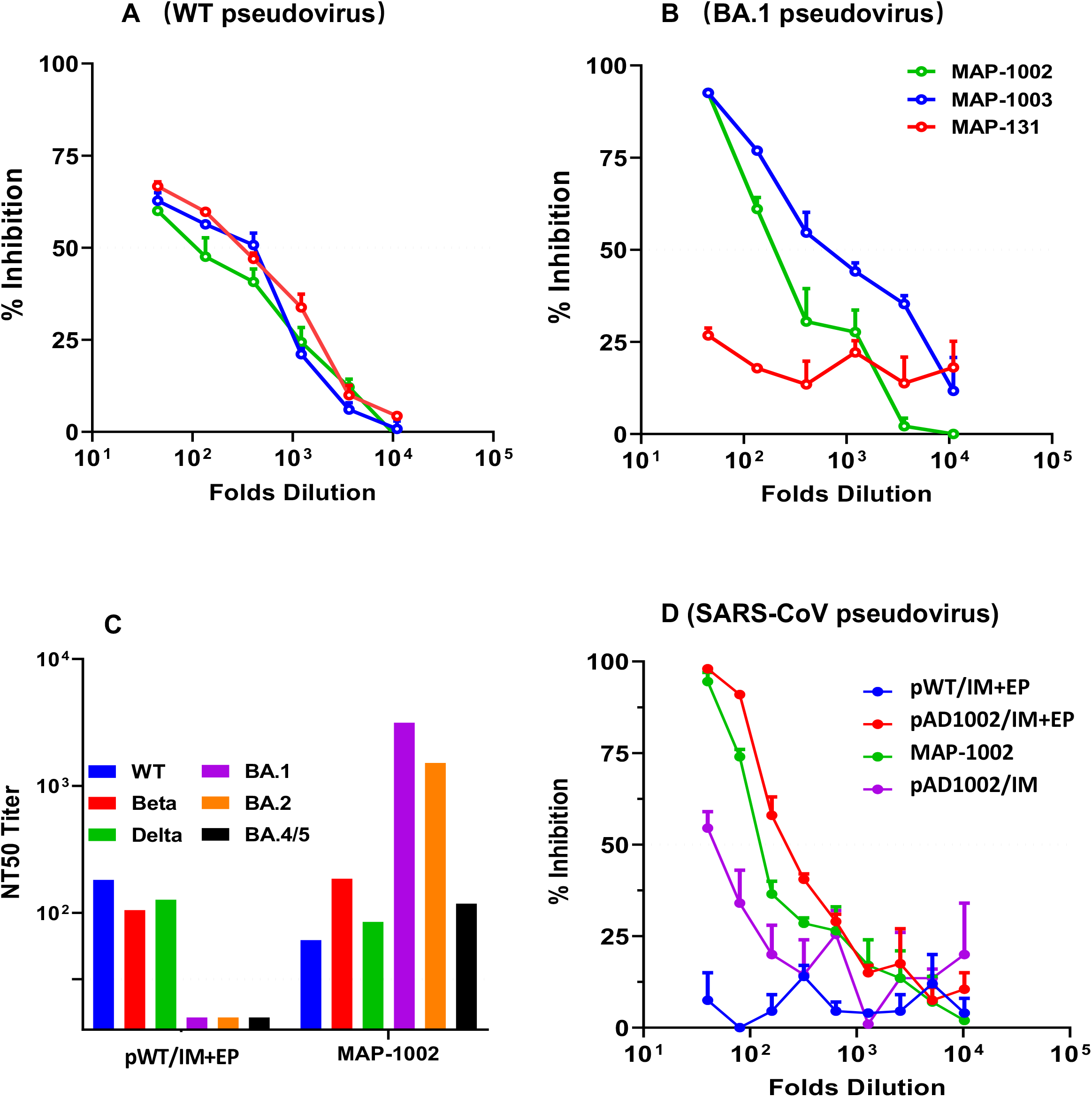
Broad spectrum neutralization of mouse antibodies induced by MAP-mediated DNA vaccination. (**A, B**) BALB/c mouse serum samples collected 14 days post boost immunization with MAP-1002, MAP-1003 and MAP-131 were tested for ability to block mimic infection of ACE2-expressing HEK293T cells by pseudo-viruses displaying S protein of SARS-CoV-2 prototypeor Omicron BA.1. The results are expressed as percentage inhibition of infection. (**C**) Antisera from BALB/c mice immunized with MAP-1002 or pWT/IM + EP were tested for neutralization of pseudo-typed virus of SARS-CoV-2 prototype, Beta, Delta, Omicron BA.1, BA.2 and BA.4/5. The values are NT50 titers. (**D**) Antisera from BALB/c mice immunized with MAP-1002, or pAD1002/IM, or pAD1002/IM + EP, or pWT/IM + EP were compared for ability to block mimic infection of ACE2-expressing HEK293T cells by SARS-CoV pseudo-virus. The horizontal dashed line indicates the limit of detection. Data are means ± SEM.

To gain further insight on the neutralization spectra of DNA vaccine-generated Abs, we next compared MAP-1002 and pWT/IM + EP antisera for ability to block mimic infection of ACE2-transgenic HEK293T cells by a set of pseudoviruses displaying S protein of WT SARS-CoV-2 or variant Beta and Delta, or Omicron subvariant BA.1, BA.2 or BA.4/5. As shown in **Fig. 6C**, MAP-1002 antisera neutralized all 6 pseudo-viruses tested, albeit BA.4/5 neutralization titer was relatively lower than that against BA.1 and BA.2. By contrast, pWT/IM + EP antisera neutralized WT, Beta and Delta, but not any of the 3 Omicron pseudo-viruses. Additionally, mouse antisera induced by MAP-1002 and pAD1002/IM + EP, but not pWT/IM + EP, immunization effectively blocked mimic infection of ACE2-transgenic HEK293T cells by pseudo-virus displaying S protein of SARS-CoV, while antisera from pAD1002/IM vaccinated animals showed only marginal blocking effect (**Fig. 6D**). These results provide proof of concept evidence that MAP^Adv^-based DNA vaccines encoding RBD^SARS^-containing chimera have the potential to provide broad-spectrum protection against SARS-CoV-2 VOCs as well as other heterologous Sarbecoviruses.

## Discussion

Effective vaccines that can provide broad coverage against existing and newly emerging variants of SARS-CoV-2 are urgently needed. Previous investigators showed that tandem RBD^WT^ homodimer or heterodimeric RBD of SARS-CoV-2 WT-Beta and Delta-BA.1 were able to generate cross-protective immunity against SARS-CoV-2 VOCs in models of mouse and rhesus monkey (16,17). Here we demonstrate DNA constructs encoding chimeric RBD containing RBD^SARS^ (pAD1002 and pADV131) are significantly more immunogenic than that encoding heterodimer RBD without RBD^SARS^ or FL S protein-encoding plasmid pWT in terms of inducing specific IgG responses in vivo (**Fig. 1**). The apparent immune-boosting effect of RBD^SARS^ should be of value in the future design of vaccines against SARS-related coronaviruses.

Immunogenicity-boosting of the DNA vaccine candidates can be significantly enhanced when intradermally delivered using MAP^Adv^. In fact, MAP^Adv^ was as effective as IM + EP in DNA vaccine delivery based on generation of RBD-specific IgG responses in model animals of different genetic backgrounds. MAP-1002, the most immunogenic amongst the three RBD-encoding vaccine candidates prepared in the present study, elicited strong and long-lasting binding and neutralizing IgG against SARS-CoV, WT SARS-CoV-2, antigen-matched as well as antigen-unmatched SARS-CoV-2 variants. Moreover, MAP-1002 significantly outperformed inactivated virus vaccine in eliciting RBD-specific IFN-γ^+^ CD8^+^ CTL cells in mice (**Fig. 5**). We believe that MAP-based heterodimeric RBD-encoding DNA constructs, as represented by MAP-1002, can serve as a potential COVID-19 vaccines for further translational studies.

In recent years, the possibility of replacing EP and gen-gun with MAPs for equally effective yet much less uncomfortable DNA vaccination has been extensively studied. Different forms of MAPs have been developed for intradermal delivery of naked DNA plasmids or nanoparticle DNA vaccines against infectious disease or cancer (29–31). Despite much progress in this field, however, molecular mechanisms for the impressive effectiveness of MAP-mediated DNA immunization are still poorly understood. In our study, strong and durable intradermal expression of the Luc gene on site of the MAP-mediated delivery was observed (**Fig. 3**). Needle-injected plasmid pVAX-Luc was also intradermally expressed, but the Luc gene expression in this case was comparatively less intense and durable (data not shown). One possible explanation for this phenomenon is that MN penetration (stimulation) accompanying MAP application triggers skin tissue cells to endocytose (and subsequently express) the unloaded DNA molecules. The possibility that excipients used for MN fabrication played a major role in facilitating the uptake and expression of MAP-delivered genes can be excluded, because i.d. injection of 20 μg pVAX1-Luc plasmid in 30 μl excipient solution or PBS produced similarly weak bioluminescence signals at the injection sites (data not shown). The skin layers contain abundant antigen presenting cells (APCs) such as LCs and dendritic cells (DCs) that play important roles in inducing adaptive immunity (26–28). Vaccine DNA unloaded from an applied 1 cm^2^ skin patch will directly reach some 100,000 APCs in dermis layer, which could uptake DNA plasmids and then migrate to draining LNs as matured APCs expressing the encoded antigen, thereby triggering strong adaptive immunological responses (35).

The successful control of COVID-19 pandemic relies not only on the development of vaccines, but also on the storage, transportation, distribution and effective administration of vaccines. Currently available inactivated virus, subunit or nucleic acid COVID-19 vaccines must be stored in either 4°C or a frozen state that may hinder their application in developing countries. By contrast, MAP-based DNA vaccines are much more stable and do not require refrigerated conditions for storage and distribution. When MAP-1002 was maintained at 25°C for one month, for example, no observable decrease of immunogenicity compared to those kept in 4°C refrigerator was found in terms of ability to induce specific IgG responses in mice (**supplemental Fig. S3**). Such stable vaccines could even be posted to rural areas for family member-assisted or self-administration under extreme circumstances.

Most current vaccines, including those approved for COVID-19, are administered i.m. or s.c. using hypodermic needles. However, there are several disadvantages for such vaccines including pain and fear of needlestick, the need for trained healthcare professionals for vaccine administration. MAPs offer a painless and minimally invasive alternative to the traditional vaccination methods by directly deposing vaccines amongst a dense population of key immune cells just below the skin surface. The administration of MAP-based vaccines is as simple as using Band-Aids and takes about 10-15 minutes to complete. Increase MAP application time from 15 to 30 or 60 min did not enhance levels of serological IgG titers of recipient animals (**supplemental Fig. S4**), suggesting 15 min was enough for DNA load in MAP^Adv^ to be delivered to the dermis tissue.

In 2021 WHO listed COVID-19 pandemic, vaccine hesitancy and limited vaccine global accessibility as top challenges to global health. By combining the heterodimeric fusion RBD approach, DNA vaccination and MAP technology, we prepared DNA vaccine candidates that can address all these challenges. This work may lay the foundation for developing promising broadly protective, thermal stable and easy operating DNA vaccines for combating COVID-19 pandemic.

## ACKNOWLEDGEMENT

This work was partially supported by a Major Research & Development Program grant from the Ministry of Science and Technology of China (2021YFC2302500). We thank Inovio for kindly providing CELLECTRA^®^2000 and needle electrode used in this study.

## AUTHOR CONTRIBUTIONS

FF, XZ, YD, XX, YZ, LW, FYG, YYP performed experiments; LJ and JK prepared the MAPs. XMG, FF, BW, GFY, ZK and GZ analyzed and interpreted data; FF, YD and FYG prepared the figures; XMG and BW provided conceptual advice and coordinated the study. XMG wrote the paper. BW, GZ and GFY reviewed and commented on the manuscript before submission.

## SUPPLEMENTAL MATERIALS

All data are available in the main text or the Supplemental information which can be found online at https://doi.org/........

