## Supplemental figures for "Potent Immunogenicity and Broad-Spectrum Protection Potential of Microneedle Array Patch-Based COVID-19 DNA Vaccine Candidates Encoding Dimeric RBD Chimera of SARS-CoV and SARS-CoV-2 Variants"

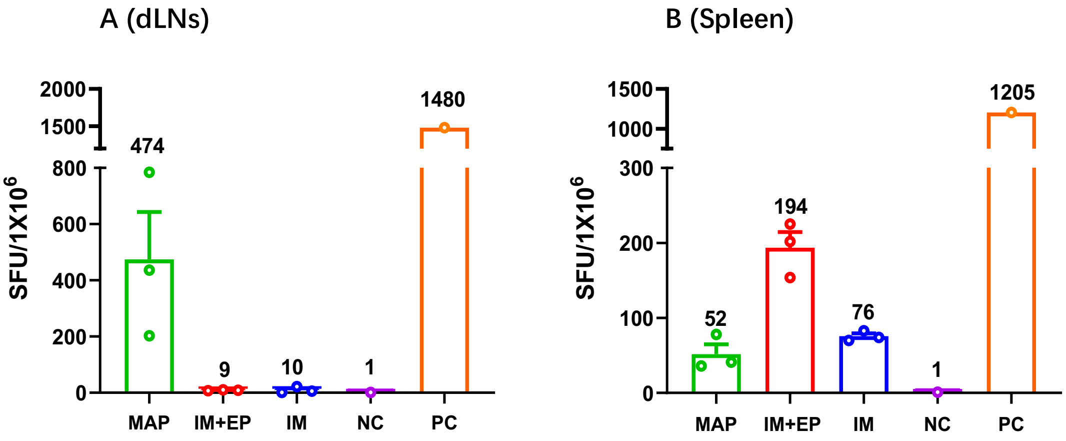


**Figure S1. RBD-specific IFN-γ$^{+}$ cell response induced by electroporated pAD1002**. Draining LN cells (**A**) and splenocytes (**B**) from BALB/c mice (n=5) that had been twice immunized with MAP-1002 (MAP), or pAD1002/IM+EP, or pAD1002/IM were used in ELISpot analysis of IFN-γ spot-forming cells (SFU) after re-stimulation with pooled 14-mer overlapping RBD$^{WT}$ peptides. LNs and spleens from unimmunized mice were included as negative control (NC). Mitomycin-stimulated splenocytes were included as positive control (PC). The results are shown as IFN-γ SFU per million cells. Data represent mean ± SD (n= 5 biologically independent samples).

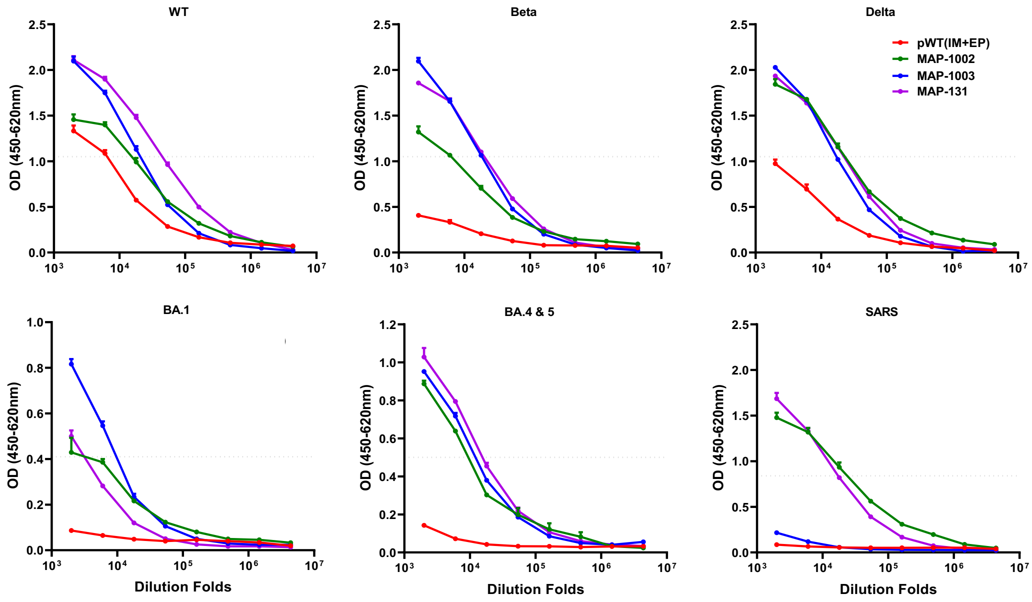


**Figure S2. Comparison of RBD-binding titers of murine serological IgG.** Serum samples from BALB/c mice (n=5 per group), collected 14 days after boost immunization with MAP-1002, MAP-1003, MAP-131 or 20 μg pWT/IM+EP, were individually titrated against recombinant RBD of WT, Beta, Delta, Omicrion BA.1, BA.4/5 of SARS-CoV-2, or SARS-CoV RBD, in ELISAs using HRP-labeled anti-mouse IgG for detection. The results (mean ± SD) are shown as OD (450-650 nm).

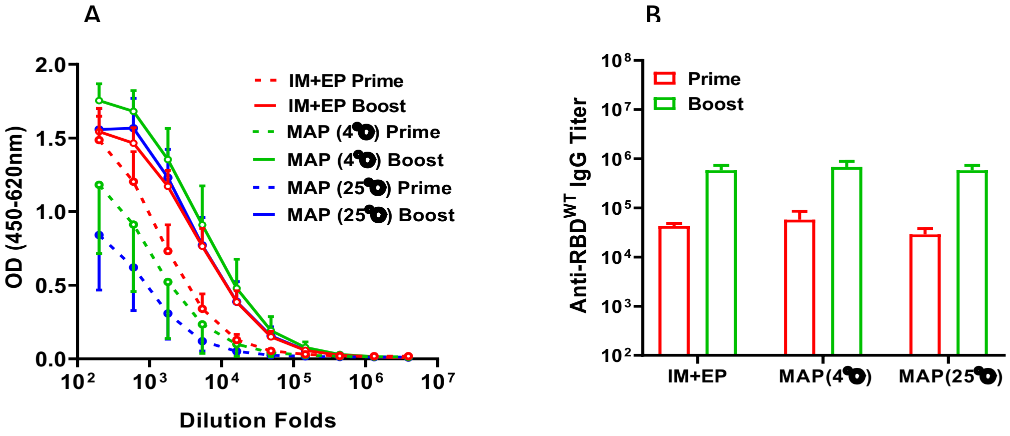


**Figure S3. Immunogenicity of MAP-1002 after storage at $4^{0}$C or $25^{0}$C.** Sample of MAP-1002 were stored at $4^{0}$C or $25^{0}$C for 30 days and then used to immunize BALB/c mice (two doses with fortnight interval). Serum samples from the vaccinated animals 14 days after prime and boost immunizations were titrated against recombinant RBD$^{WT}$ in ELISA using HRP-labeled anti-mouse IgG for detection (n=5 biologically independent samples). Sera from mice vaccinated with 20 μg pAD1002/IM+EP (two doses with fortnight interval) were included as controls. Data represent mean ± SD of (**A**) OD$_{450-650nm}$ and (**B**) calculated endpoint dilution titers of IgG.

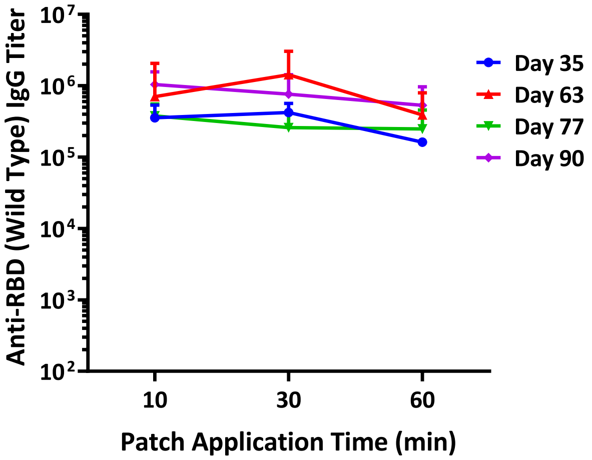


**Figure S4. IgG responses in BALB/c mice induced by MAP-131 administration.** MAP-131 patches were applied to the shaved skin surface of BALB/c mice with thumb pressure and allowed to stay for 15, 30 or 60 min before removal. Serum samples from the vaccinated animals, collected on days 35, 63, 77 and 90 after primary immunization, were titrated against recombinant $RBD^{WT}$ in ELISA. The results are expressed as endpoint dilution titers of $RBD^{WT}$-specific serological IgG. Data represent mean ± SD (n= 5 biologically independent samples).
